## Supplementary Figures for "Transcription Termination and Antitermination of Bacterial CRISPR Arrays"

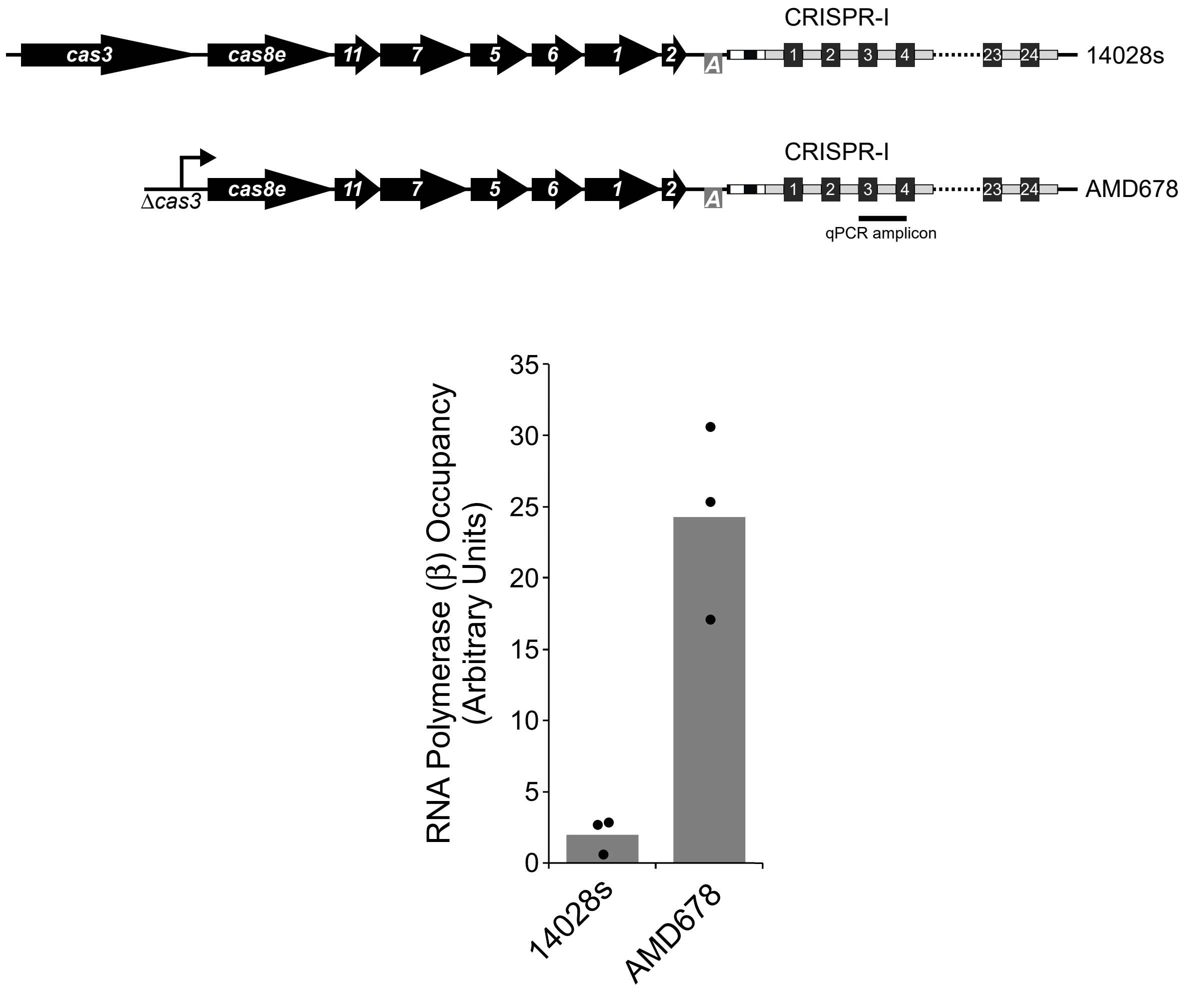


**Figure 1 - figure supplement 1. The CRISPR-I array is co-transcribed with the upstream *cas* gene operon.** Occupancy of RNA polymerase (β subunit) measured by ChIP-qPCR within the CRISPR-I array in wild-type 14028s or AMD678, which has a constitutive promoter in place of *cas3*, as indicated in the schematic. The location of the qPCR amplicon is indicated on the schematic. Values shown with bars are the average of three independent biological replicates, with dots showing each individual datapoint.

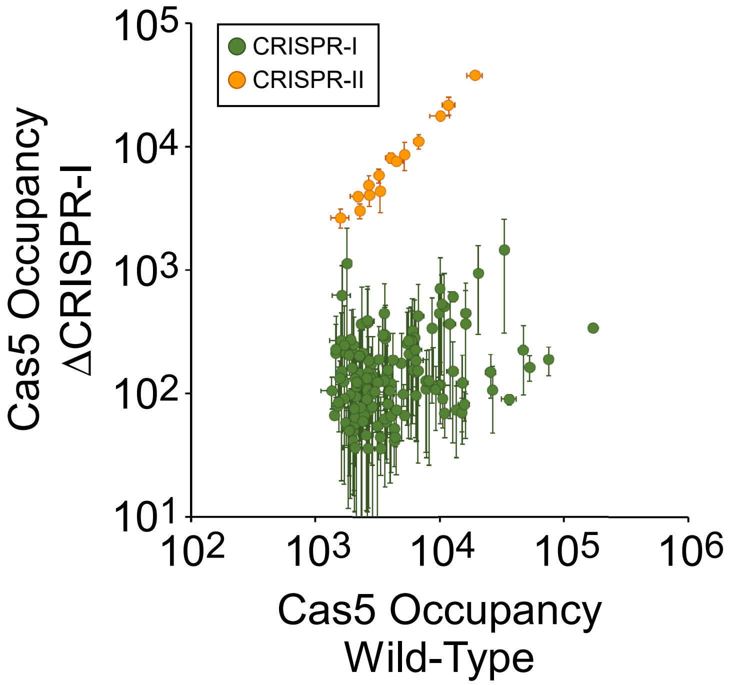


**Figure 3 - figure supplement 1. Confirmation of spacer assignments to Cas5 binding sites.** Comparison of $FLAG_3$-Cas5 ChIP-seq occupancy at off-target chromosomal sites in cells with an intact CRISPR-I array (AMD678; *x*-axis), and ΔCRISPR-I cells (AMD679; *y*-axis). Cascade binding associated with spacers from CRISPR-I is indicated by green datapoints. Cascade binding associated with spacers from CRISPR-II is indicated by orange datapoints. Values plotted are the average of two independent biological replicates. Error bars represent one standard-deviation from the mean.

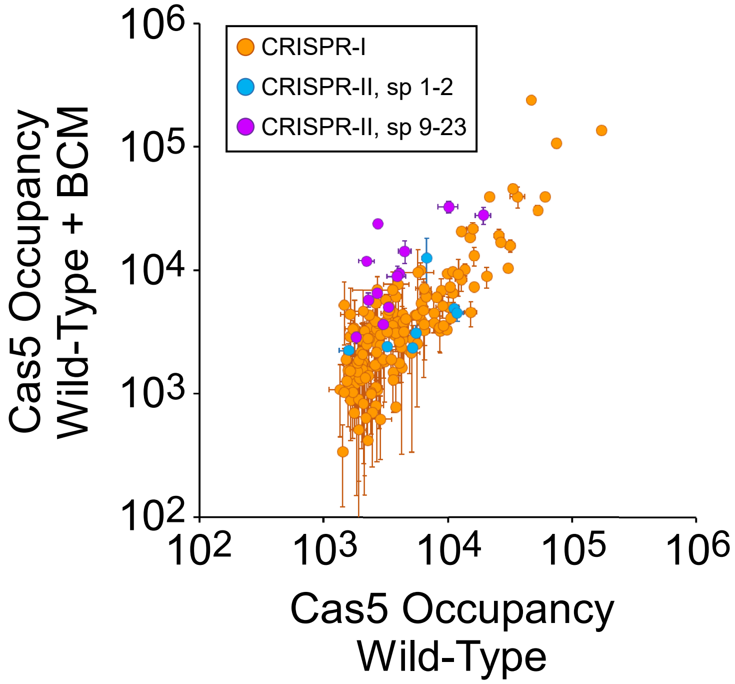


**Figure 4 - figure supplement 1. Inhibition of Rho in cells with an intact boxA increases the use of spacers 9-23 of CRISPR-II.** Comparison of $FLAG_3$-Cas5 ChIP-seq occupancy at off-target chromosomal sites in cells with an intact CRISPR-II *boxA* (AMD678) without (*x*-axis) and with bicyclomycin (BCM) treatment (*y*-axis). Cascade binding associated with spacers from CRISPR-I is indicated by orange datapoints. Cascade binding associated with spacers 1-2 from CRISPR-II is indicated by light blue datapoints. Cascade binding associated with spacers 9-23 from CRISPR-II is indicated by purple datapoints. Values plotted are the average of two independent biological replicates. Error bars represent one standard-deviation from the mean.

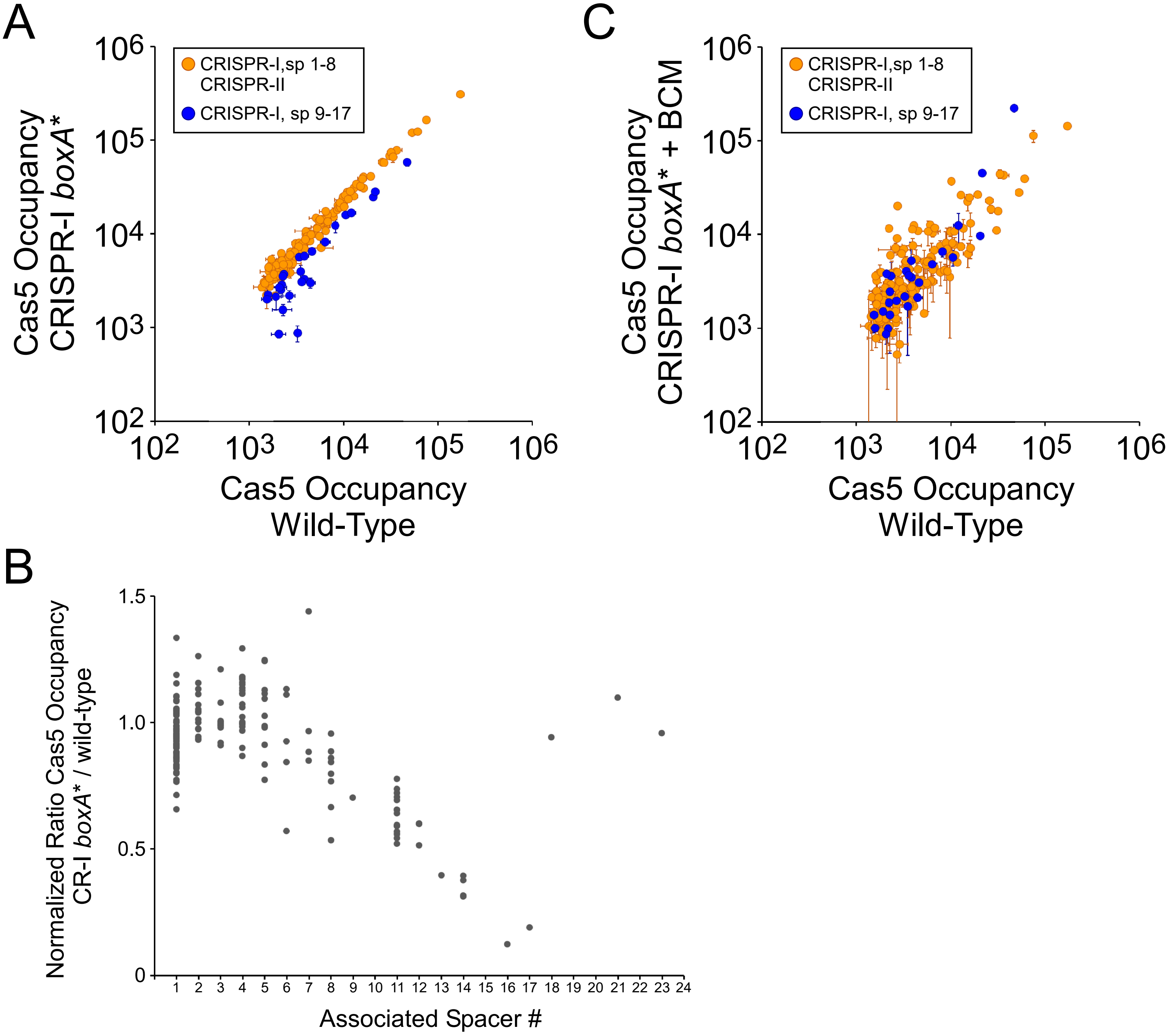


**Figure 4 - figure supplement 2. The CRISPR-I *boxA* facilitates use of all spacers by preventing premature Rho termination. (A)** Comparison of $FLAG_3$-Cas5 ChIP-seq occupancy at off-target chromosomal sites in cells with an intact CRISPR-I *boxA* (AMD678; *x*-axis), and cells with a single base-pair substitution in the CRISPR-I boxA ("*boxA**"; AMD684; *y*-axis). Cascade binding associated with spacers from CRISPR-II, and spacers 1-8 from CRISPR-I, is indicated by orange datapoints. Cascade binding associated with spacers 9-17 from CRISPR-I is indicated by dark blue datapoints. Values plotted are the average of two independent biological replicates. Error bars represent one standard-deviation from the mean. **(B)** Normalized ratio of $FLAG_3$-Cas5 occupancy associated with spacers from CRISPR-I. Values are plotted according to the associated spacer, and are normalized to the average value for sites associated with spacers from CRISPR-II. **(C)** Comparison of $FLAG_3$-Cas5 ChIP-seq occupancy at off-target chromosomal sites in cells with an intact CRISPR-I boxA (AMD678; *x*-axis), and cells with a single base-pair substitution in the CRISPR-I boxA ("*boxA**"; AMD684) that were treated with bicyclomycin (BCM; *y*-axis).

CAATATATGGGTTTATTACTGTGCTCTTTAACAATATATTGGTGTCTTCCCCACGCAGGTGGGGGTGTTTCTGA
TAAAACTTACGAATTGTTTATTAGCGATGGGTCTTCCCCACGCAGGTGGGGGTGTTTCTATAGAGGATCACCAT
AATATTAACGTAAAAATGTCTTCCCCACGCAGGTGGGGGTGTTTCCACTGACTCGCCAAGCTTCGCCACCGCTT
CTAGGTCTTCCCCACGCAGGTGGGGGTGTTTCCACCGCTAATCATGGTGGAACGAACGCCATCAAGTCTTCCCC
ACGCAGGTGGGGGTGTTTCTGATTTTGGAAGTAATGGGAACTGAGCGTTAAGGTCTTCCCCACGCAGGTGGGGG
TGTTTCCCAAAAACCTACGCGGTTTTAAATGGATTCGACGTCTTCCCCACGCAGGTGGGGGTGTTTCTATCTTG
GTTTTGCAGGTTGTTAATCTCAGCGTGTCTTCCCCACGCAGGTGGGGGTGTTTCTGATTGGTTCCAGTTTATGA
CAAGAACCAACACGTCTTCCCCACGCAGGTGGGGGTGTTTCCGACTTTTGCATCATCGATGTACGGAACGCTAG
GTCTTCCCCACGCAGGTGGGGGTGTTTCCACTGAGATTGCGTGTCGCCGACTTGCGCTTGCGTCTTCCCCACGC
AGGTGGGGGTGTTTCTAGACTATCAATGTGCGCTTGCAAGTCTTTTAAGTCTTCCCCACGCAGGTGGGGGTGTT
TCTTGATCGCTCTGAAATTGTGACTTGTTTTGTTAGTCTTCCCCACGCAGGTGGGGGTGTTTCTAGATCTTAAT
TGTTCGCGTTGAATGGAAATTGTCTTCCCCACGCAGGTGGGGGTGTTTCCAGATTGTAGATAAGCAGGAGACT
GCCCACCAGGTCTTCCCCACGCAGGTGGGGGTGTTTCCGCTTCTGTAGAGGTGATGGGTCCAAAGATGTTGTCT
TCCCCACGCAGGTGGGGGTGTTTCTACATTCGTGATATCAGCGGATGCGCTTGGTCAGTCTTCCCCACGCAGGT
GGGGGTGTTTCCAAGGAAATTTGCTACCAAAAGACGTGATCGAGGTCTTCCCCACGCAGGTGGGGGTGTTTCCC
AAGCAGCAATAAAAATGGAGAGCAATCCTATGTCTTCCCCACGCAGGTGGGGGTGTTTCCCAAAGCAATTCAAC
AGAGGCCATCAATCGCCTGTCTTCCCCACGCAGGTGGGGGTGTTTCTATTGCATTGAATTGCCCCACTCTTTGA
CCACAGTCTTCCCCACGCAGGTGGGGGTGTTTCTCAAAATACTTCTCGCACATCCTGCAAGCGCTGGTCTTCCC
CACGCAGGTGGGGGTGTTTCCCGAATACAACGGGCAAGCCTGCGCTAACGTTCGTCTTCCCCACGCAGGTGGGG
GTGTTTCCGGTTGATAAAACGCTGCGTAAGTTTTTCGAAGGTCTTCCCCACGCAGGTGGGGGTGTTTCTAAAAC
TTCATAGATTGTTGCCTCCATTGTTTCGTCTTCCCCACGCAGGTGGGGGTGTTTCTACGTCAACGGACAAACCA
AAACCGAATGGAAGGTCTTCCCCACGCAGGTGGGGGTGTTTCCACTGAGATTGCGTGTCGCCGACTTGCGCTTG
CGTCTTCCCCACGCAGGTGGGGGTGTTTCTACTGGCCGATGAGGTGGACCGCTACGGCTTCAGTCTTCCCCACG
CAGGTGGGGGTGTTTCTCACCCAGCACATTACCACCCATGATCAGCGTTGTCTTCCCCACGCAGGTGGGGGTGT
TTCCGAGTGGGTTTAGGTTGTAGGTTGCACATACGCGTCTTCCCCACGCAGGTGGGGGTGTTTCTTCGAAAAGC
TATTAGGCGGCATAACCACAGTTGTCTTCCCCACGCAGGTGGGGGTGTTTCCAACAAACCAGCCACTTTGCATT
TTGTAGCAGAGTCTTCCCCACGCAGGTGGGGGTGTTTCTGGAAGTTATTTATATTGGACCTGATTGCACGGGTC
TTCCCCACGCAGGTGGGGGTGTTTCTACTTTGGGCCTTGTTTTATGTGCCAGTGCCCGGTCTTCCCCACGCAGG
TGGGGGTGTTTCCCACTATTTCAATAATCGGCGTAGCTCCGACTGGTCTTCCCCACGCAGGTGGGGGTGTTTCC
GTAATCTTCTTTGTCTGAGTAATCCAAAATACGTCTTCCCCACGCAGGTGGGGGTGTTTCTCATGGCAGCGATA
TTTGTTTTACCCTTCATACGTCTTCCCCACGCAGGTGGGGGTGTTTCCATCTGGAACGCACTCAAGCGGCAATC
CGAAGTGTCTTCCCCACGCAGGTGGGGGTGTTTCTACGTCCAGCATTACCGCCGCGCCGTGTCGAGTGTCTTCC
CCACGCAGGTGGGGGTGTTTCTCCTCCCTGCTCATATGCCGTTAAAACTTTCTCGTCTTCCCCACGCAGGTGGG
GGTGTTTCT

Spacer 1 GATAAAACTTACGAATTGTTTATTAGCGATGG
Spacer 2 ATAGAGGATCACCATAATATTAACGTAAAAAT
Spacer 3 ACTGACTCGCCAAGCTTCGCCACCGCTTCTAG
Spacer 4 ACCGCTAATCATGGTGGAACGAACGCCATCAA
Spacer 5 GATTTTGGAAGTAATGGGAACTGAGCGTTAAG
Spacer 6 CAAAAACCTACGCGGTTTTAAATGGATTCGAC
Spacer 7 ATCTTGGTTTTGCAGGTTGTTAATCTCAGCGT
Spacer 8 GATTGGTTCCAGTTTATGACAAGAACCAACAC
Spacer 9 GACTTTTGCATCATCGATGTACGGAACGCTAG
Spacer 10 ACTGAGATTGCGTGTCGCCGACTTGCGCTTGC
Spacer 11 AGACTATCAATGTGCGCTTGCAAGTCTTTTAA
Spacer 12 TGATCGCTCTGAAATTGTGACTTGTTTTGTTA
Spacer 13 AGATCTTAATTGTTCGCGTTGAATGGAAATT
Spacer 14 AGATTGTAGATAAGCAGGAGACTGCCCACCAG
Spacer 15 GCTTCTGTAGAGGTGATGGGTCCAAAGATGTT
Spacer 16 ACATTCGTGATATCAGCGGATGCGCTTGGTCA

```
Spacer 17 AAGGAAATTTGCTACCAAAAGACGTGATCGAG
Spacer 18 CAAGCAGCAATAAAAATGGAGAGCAATCCTAT
Spacer 19 CAAAGCAATTCAACAGAGGCCATCAATCGCCT
Spacer 20 ATTGCATTGAATTGCCCCACTCTTTGACCACA
Spacer 21 CAAAATACTTCTCGCACATCCTGCAAGCGCTG
Spacer 22 CGAATACAACGGGCAAGCCTGCGCTAACGTTC
Spacer 23 GGTTGATAAAACGCTGCGTAAGTTTTTCGAAG
Spacer 24 AAAACTTCATAGATTGTTGCCTCCATTGTTTC
Spacer 25 ACGTCAACGGACAAACCAAAACCGAATGGAAG
Spacer 26 ACTGAGATTGCGTGTCGCCGACTTGCGCTTGC
Spacer 27 ACTGGCCGATGAGGTGGACCGCTACGGCTTCA
Spacer 28 CACCCAGCACATTACCACCCATGATCAGCGTT
Spacer 29 GAGTGGGTTTAGGTTGTAGGTTGCACATACGC
Spacer 30 TCGAAAAGCTATTAGGCGGCATAACCACAGTT
Spacer 31 AACAAACCAGCCACTTTGCATTTTGTAGCAGA
Spacer 32 GGAAGTTATTTATATTGGACCTGATTGCACGG
Spacer 33 ACTTTGGGCCTTGTTTTATGTGCCAGTGCCCG
Spacer 34 CACTATTTCAATAATCGGCGTAGCTCCGACTG
Spacer 35 GTAATCTTCTTTGTCTGAGTAATCCAAAATAC
Spacer 36 CATGGCAGCGATATTTGTTTTACCCTTCATAC
Spacer 37 ATCTGGAACGCACTCAAGCGGCAATCCGAAGT
Spacer 38 ACGTCCAGCATTACCGCCGCGCCGTGTCGAGT
Spacer 39 CCTCCCTGCTCATATGCCGTTAAAACTTTCTC
```

**Figure 6 – figure supplement 1. Sequence of the CRISPR array and upstream sequence from *Vibrio cholerae* A50.** The first nucleotide shown is the first nucleotide downstream of *cas2*. Repeat sequences are highlighted in green. The *boxA* sequence is highlighted in yellow. Individual spacers and their position in the array is indicated below the sequence of the array. The red highlighted spacers are identical.

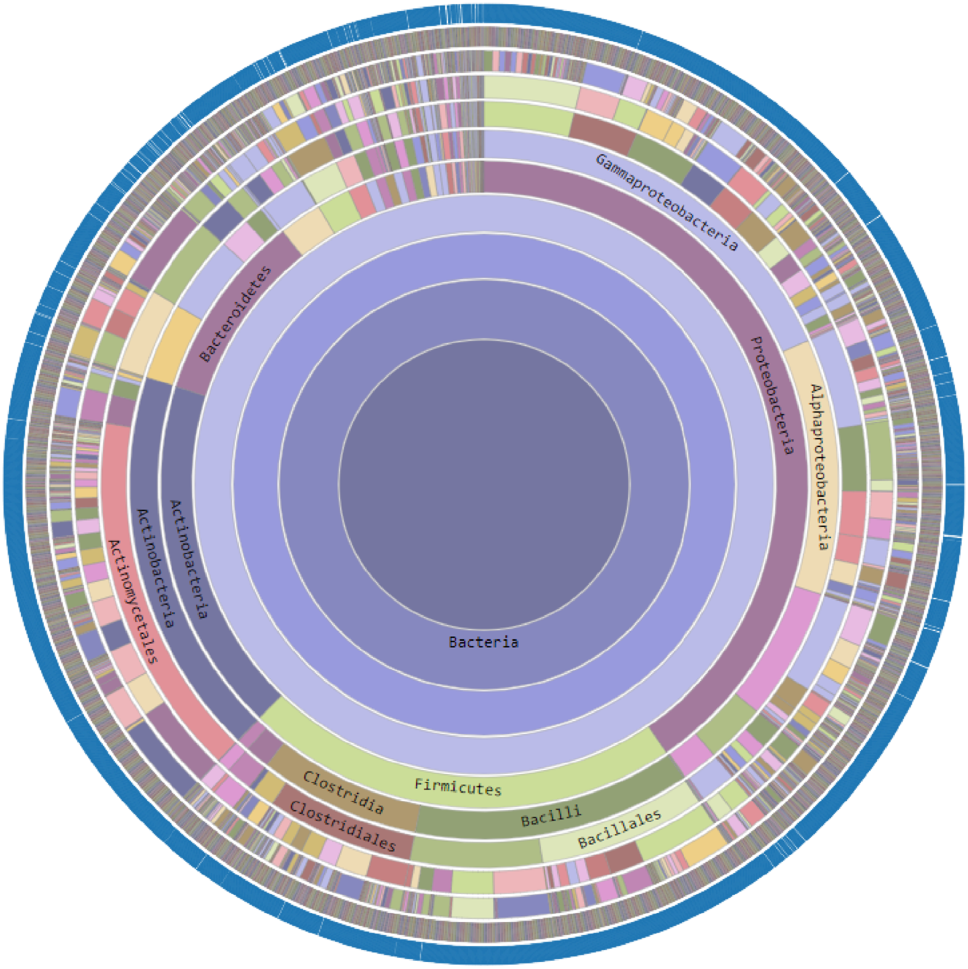


**Figure 7 - figure supplement 1. Conservation of *nusB* across the bacterial kingdom.** A conservation plot generated using the Aquerium tool (Adebali and Zhulin, 2017). Each circle represents a taxonomic rank (kingdom, phylum, class, order, family, genus, species), with lower level ranks being further from the center of the plot. The outer, blue circle indicates species that have *nusB*. Raw data are shown in Table S3.

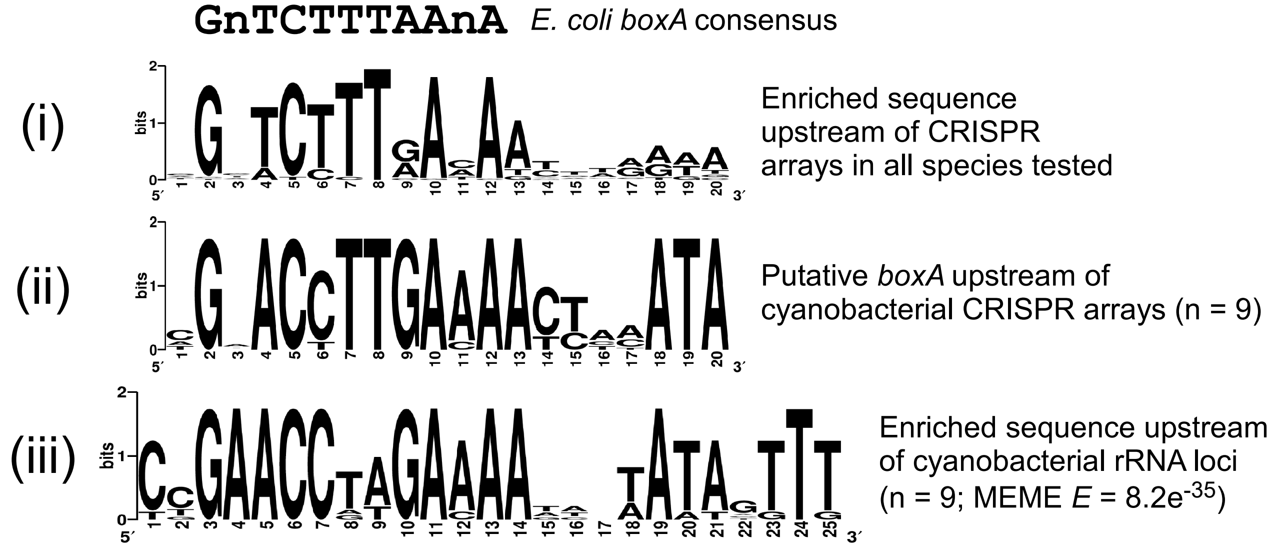


**Figure 7 - figure supplement 2. Analysis of putative *boxA* sequences upstream of cyanobacterial CRISPR arrays and ribosomal RNA loci.** Comparison of sequence motifs for putative *boxA* sequences from (i) the set of CRISPR array upstream sequences from 187 bacterial species (also shown in Figure 7A), (ii) the subset of (i) from 9 cyanobacterial species, and (iii) ribosomal RNA (rRNA) upstream sequences for the 9 species analyzed in (ii). The *E*-value determined by MEME is shown for the motif derived from cyanobacterial rRNA upstream sequences. The E. coli *boxA* consensus sequence is shown above the sequence logos.
